# Cellular mechanisms of chick limb bud morphogenesis

**DOI:** 10.1101/2020.09.10.292359

**Authors:** Gaja Lesnicar-Pucko, Julio M Belmonte, Marco Musy, James A. Glazier, James Sharpe

**Affiliations:** Centre for Genomic Regulation (CRG), Barcelona Institute of Science and Technology (BIST), Dr. Aiguader 88, 08003 Barcelona, Spain; Universitat Pompeu Fabra (UPF), Dr. Aiguader 88, 08003 Barcelona, Spain; Biocomplexity Institute and Department of Physics, Indiana University Bloomington, Bloomington, Indiana, 47405, USA; European Molecular Biology Laboratory (EMBL) Heidelberg, Developmental Biology Unit, Meyerhofstraße 1, Heidelberg 69117 Germany; Physics Department, North Carolina State University, 2401 Stinson Drive, Raleigh, NC, USA; European Molecular Biology Laboratory (EMBL) Barcelona, Dr. Aiguader 88, 08003 Barcelona, Spain; Institucio’ Catalana de Recerca i Estudis Avancats (ICREA), Pg. Lluís Companys 23, 08010 Barcelona, Spain

**Keywords:** morphogenesis, Cellular Potts Model, *in ovo*, intercalation, limb bud, 2-photon microscopy

## Abstract

Although some of the molecular pathways involved in limb bud morphogenesis have been identified, the cellular basis of the process is not yet understood. Proposed cell behaviours include active cell migration and oriented cell division, but ultimately, these questions can only be resolved by watching individual mesenchymal cells within a completely normal developmental context. We developed a minimally-invasive *in ovo* two-photon technique, to capture high quality time-lapse sequences up to 100 microns deep in the unperturbed growing chick limb bud. Using this technique, we characterized cell shapes and other oriented behaviours throughout the limb bud, and found that cell intercalation drives tissue movements, rather than oriented cell divisions or migration. We then developed a 3D cell-based computer simulation of morphogenesis, in which cellular extensions physically pull cells towards each other, with directional bias controlled by molecular gradients from the ectoderm (Wnts) and the Apical Ectodermal Ridge (FGFs). We defined the initial and target shapes of the chick limb bud in 3D by OPT scanning, and explored which orientations of mesenchymal intercalation correctly explain limb morphogenesis. The model made a couple of predictions: Firstly, that elongation can only be explained when cells intercalate along the direction towards the nearest ectoderm. This produces a general convergence of tissue towards the central proximo-distal (PD) axis of the limb, and a resultant extension of the tissue along the PD axis. Secondly, the correct *in silico* morphology can only be achieved if the contractile forces of mesenchymal cells in the very distal region (under the Apical Ectodermal Ridge) have shorter life times than in the rest of the limb bud, effectively making the tissue more fluid by augmenting the rate of cell rearrangement. We argue that this less-organised region of mesenchyme is necessary to prevent PD-oriented intercalation events in the distal tip that would otherwise inhibit outgrowth.

## Introduction

Although the limb bud is a classical model system for organogenesis, it has mostly been used to study molecular pattern formation and spatial control of differentiation, rather than physical morphogenesis. For many years the cellular mechanism by which the limb bud achieves distal elongation was considered rather simple, and was dominated by the *proliferation gradient hypothesis* (Figure 1A) – proposing that higher proliferation rates in the distal end of the bud were largely sufficient to explain elongation (Ede and Law 1969, Reiter and Solursh 1982, Niswander and Martin 1993). We previously tested this hypothesis in a computer model utilizing quantitative empirical data, and revealed the insufficiency of this mechanism to explain the strong anisotropy of growth (Boehm, Westerberg et al. 2010). This suggested that cell behaviours in the limb bud mesenchyme are likely to be oriented (see Figure 1 for a few possible mechanisms).

**Figure 1.**
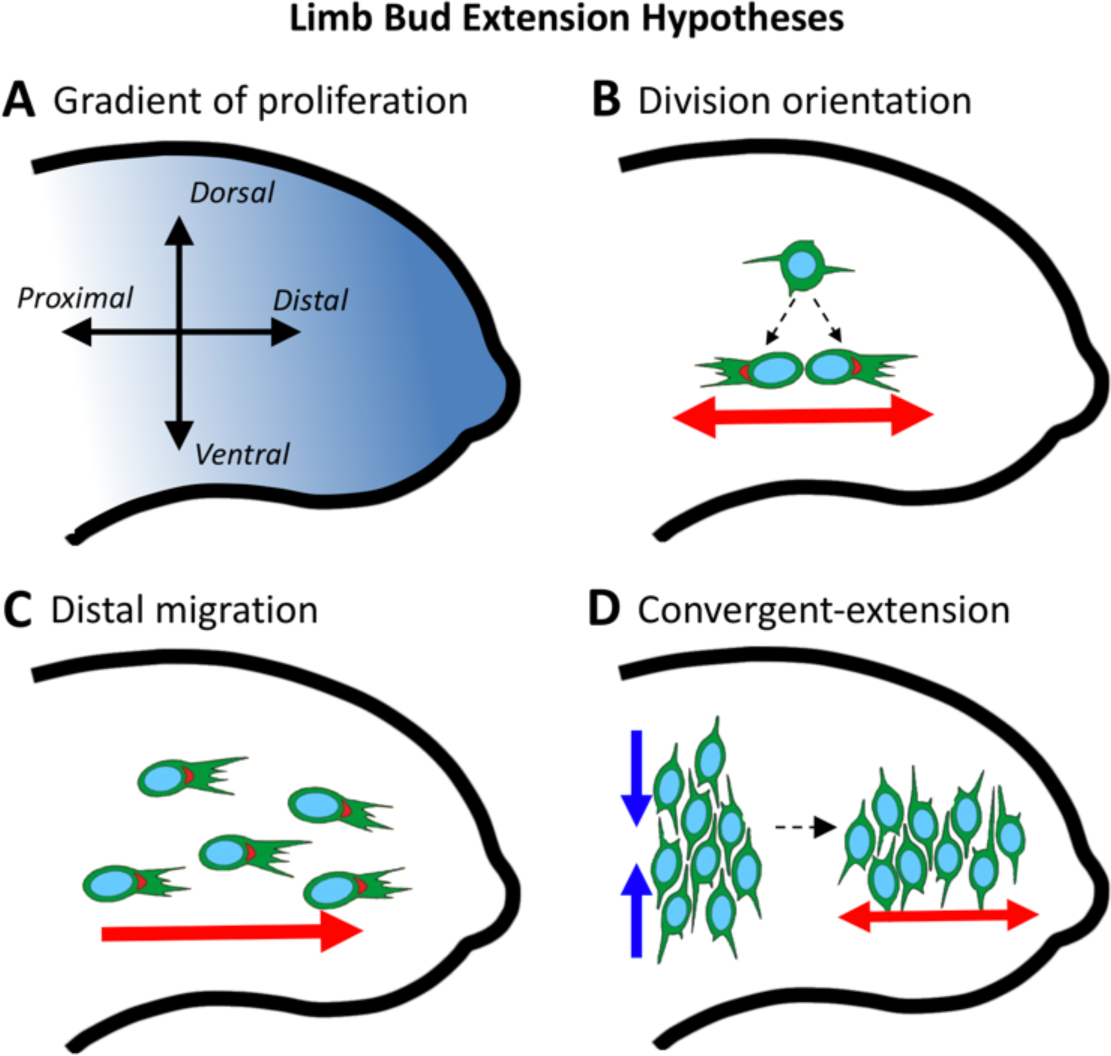
Possible cell-based mechanisms for limb bud extension. (A) *Gradient of proliferation* hypothesis. (B) *Orientation of cell division* hypothesis. (C) *Distalward migration* hypothesis. (D) *Convergent-extension* hypothesis, in which cells intercalate in the direction of the blue arrows, causing the tissue to expand in the direction of the red arrow. (A-D) Proximal-Distal (PD) axis is horizontal and Dorsal-Ventral (DV) axis is perpendicular. Anterior-Posterior axis is perpendicular to the page.

We and others, subsequently found multiple signs of oriented cellular behaviour: the cell shapes and positioning of Golgi show biases in orientation, and activities such as cell division and general cell movement are also preferentially orientated relative to the limb axes (Boehm, Westerberg et al. 2010, Gros, Hu et al. 2010, Sato, Seki et al. 2010, Wyngaarden, Vogeli et al. 2010, Gao, Song et al. 2011). The molecular basis of these behaviours has also been explored, and strong evidence exists that Wnt signalling and the PCP system are required for correct orientation of cellular activities. However, precisely which cell behaviours these molecules regulate and their relative contributions and interactions, remains unclear. For example, FGF signalling has been implicated in multiple different theories, each of which proposes a different functional role for FGF: (i) as a mitogen (Reiter and Solursh 1982), (ii) a chemoattractant for directed migration (Li and Muneoka 1999) or (iii) as a modulator of cellular motility (Benazeraf, Francois et al. 2010, Gros, Hu et al. 2010). Therefore, despite the strong evidence about which molecules are important, it is not clear which cellular processes are present during limb morphogenesis, and how much they contribute to PD elongation.

We can summarise the remaining questions as follows:

Firstly, does oriented cell division contribute to PD elongation (Figure 1B)? In *Drosophila* and zebrafish, evidence exists that the control of cell division orientation is a mechanism driving anisotropic tissue elongation (Baena-Lopez, Baonza et al. 2005, Tawk, Araya et al. 2007). At early budding stages in the mouse, knock-out of Wnt5a reduces the prevalence of distally-oriented cell divisions and impairs elongation (Gros, Hu et al. 2010), suggesting a causal relation between the two events. At later elongation stages however, the role of oriented cell division is more ambiguous. In early chick bud (Hamburger Hamilton (HH) 19) cell division orientation is biased parallel to the PD axis, compatible with PD limb bud elongation (Gros, Hu et al. 2010, Wyngaarden, Vogeli et al. 2010). Yet, at slightly later stages (HH21) cell orientation and cell divisions in the dorsal and ventral regions of the bud have been reported to be biased at 90 degrees to the PD axis, approximately along the dorso-ventral (DV) axis (Boehm, Westerberg et al. 2010, Gros, Hu et al. 2010), which would appear to work directly against PD elongation. At stage HH26, nonetheless, Sato and co-workers proposed that the bias in cell division orientation along the AP axis drives the widening of the handplate (Sato, Seki et al. 2010). In general, then, are oriented cell divisions promoting or hindering PD elongation?

Secondly, what is the contribution of active cell migration to limb elongation (Figure 1C)? Many limb mesenchymal cells appear to display a cellular orientation, *i*.*e*. they have an elongated shape, or have preferential orientation of cellular extensions. At young stages of development (HH19) the average cell orientation is distally-directed (Gros, Hu et al. 2010), suggesting that cells could theoretically migrate in the direction of bud elongation. Mesenchymal cells have also been shown to exhibit active directional chemotaxis toward an experimental source of FGF, both *in vitro* (Kim, Kang et al. 2009) and *in vivo* (Li and Muneoka 1999). It has therefore been proposed that the distal-ward migration could be controlled by a diffusible FGF, which is secreted by the Apical Ectodermal Ridge (AER) at the distal tip of the bud (Li and Muneoka 1999). However, the reported PD orientation of cells results from averaging cells across a wide region of the bud (Gros, Hu et al. 2010). Analysis of different regions separately shows instead that cells in the dorsal and ventral regions of the bud are not oriented along the PD axis, and some even appear to “migrate” directly towards and away from the nearest ectoderm (Gros, Hu et al. 2010).

The third possibility would be that mesenchymal cells might be actively intercalating to drive a convergence-extension movement (Figure 1D), similar to that involved in axis elongation during *Xenopus* and chick gastrulation (Keller, Davidson et al. 2000, Voiculescu, Bertocchini et al. 2007, Keller, Shook et al. 2008, Voiculescu, Bodenstein et al. 2014). A convergent-extension mechanism for bud elongation requires a dramatically different cellular orientation than in the cases of pure distal-migratory hypothesis or oriented cell divisions. Rather than parallel to the PD axis, cell orientations would have to be more-or-less perpendicular to the primary PD axis, reflecting the direction of intercalation (Figure 1D). The previous observations of cell orientation along the DV direction in the proximal regions may therefore support the idea of a convergent extension mechanism (Boehm, Westerberg et al. 2010, Gros, Hu et al. 2010). However, cells in the distal tip of the bud are not DV oriented.

Resolving these questions requires focusing on cellular activities, rather than molecular signalling pathways. Indeed, the limiting factor to our understanding is a lack of basic information about individual cellular behaviours (dynamic shapes, orientations and movements) during normal limb bud development. This is made more challenging by the 3D nature of the limb bud mesenchyme. While cell migration and convergent-extension have been well-studied in 2D cases, the relationships between individual cellular activities and the resulting tissue movements have not received much attention (Honda, Nagai et al. 2008, Belmonte, Swat et al. 2016). Vertebrate organogenesis is generally very sensitive to perturbations, meaning it is difficult to be sure whether cell behaviours remain developmentally normal *in vitro*.

To circumvent these difficulties, we developed an *in ovo* two-photon time-lapse approach that allows us to watch the active dynamics of individual mesenchymal cells during normal limb development in chick embryos. In contrast to slice cultures, this technique is sufficiently non-invasive that normal limb development can proceed after the imaging is completed.

In this study we first use the *in ovo* imaging technique to determine the main features and orientations of active cell movements (including cell division, active cell migration and local convergent-extension movements). We then examine the global arrangement of cellular orientations in sectioned limb bud tissues, and precisely map them throughout the limb bud. Thirdly, we develop a dynamic 3D virtual tissue simulation to explore how the observed single-cell behaviours combine to produce observed limb bud elongation and shape dynamics. These simulations predict that observed limb-bud dynamics requires a specific pattern of oriented cell intercalation in the direction towards the nearest ectoderm, and reduced cell intercalation in the distal tip of the limb bud. We suggest that FGF signalling, in addition to possibly stimulating proliferation (Niswander and Martin 1993), acting as a mitogen (Reiter and Solursh 1982), or creating a motility gradient (Li, Anderson et al. 1996) (Li and Muneoka 1999), may contribute to limb elongation by facilitating tissue stress-relaxation in the distal mesenchyme by promoting cell motility.

## Results

### A New Non-Disruptive Time-Lapse Approach to Track Mesenchymal Cells *in ovo*

*In vitro* techniques have been used to image cellular behaviour during early limb budding, when growth is still not heavily dependent on blood flow (Fallon and Todt 1984, Wanek, Muneoka et al. 1989, Gros, Hu et al. 2010, Wyngaarden, Vogeli et al. 2010). Yet, studying cell behaviours during limb elongation between stages 21HH and 23HH, requires an *in vivo* technique which does not disrupt the normal vascular supply of oxygen and nutrients. *In ovo* imaging of younger embryos has been developed previously to image neural crest migration (Kulesa and Fraser 2000), but has never been employed at later stages because the embryo moves considerably, especially due to the increasingly strong heart-beat.

We successfully adapted the *in ovo* imaging technique to later stage embryos and stabilized them enough for deep, sub-cellular resolution two-photon time-lapse imaging, without influencing normal development. To track individual cells and to reveal their shapes, we electroporated stage HH15 embryos with membrane-targeted gpiEGFP that produced a salt-and-pepper distribution of labelled cells. 24 hours after electroporation, we sealed the eggs with a plastic ring covered with a Teflon membrane, which allows normal gas exchange during imaging (Figure 2A). To dampen external vibrations and temperature changes, we submerged the egg in a beaker filled with 1% agarose, and then wrapped the beaker in a flexible heating plate (Figure 2B). To eliminate imaging artefacts due to the large movements caused by the beating heart, we stopped the heart every hour for a few minutes by cooling the embryo with of ice-cold PBS applied on top of the Teflon membrane (Figure 2C). After capturing a 3D image, we then rewarmed the embryo with warm PBS (37OC) and the heart spontaneously started to beat again. We repeated this process every hour for up to twelve hours. We also developed a post-processing program to register the images to compensate for any remaining movement that occurred during imaging.

**Figure 2.**
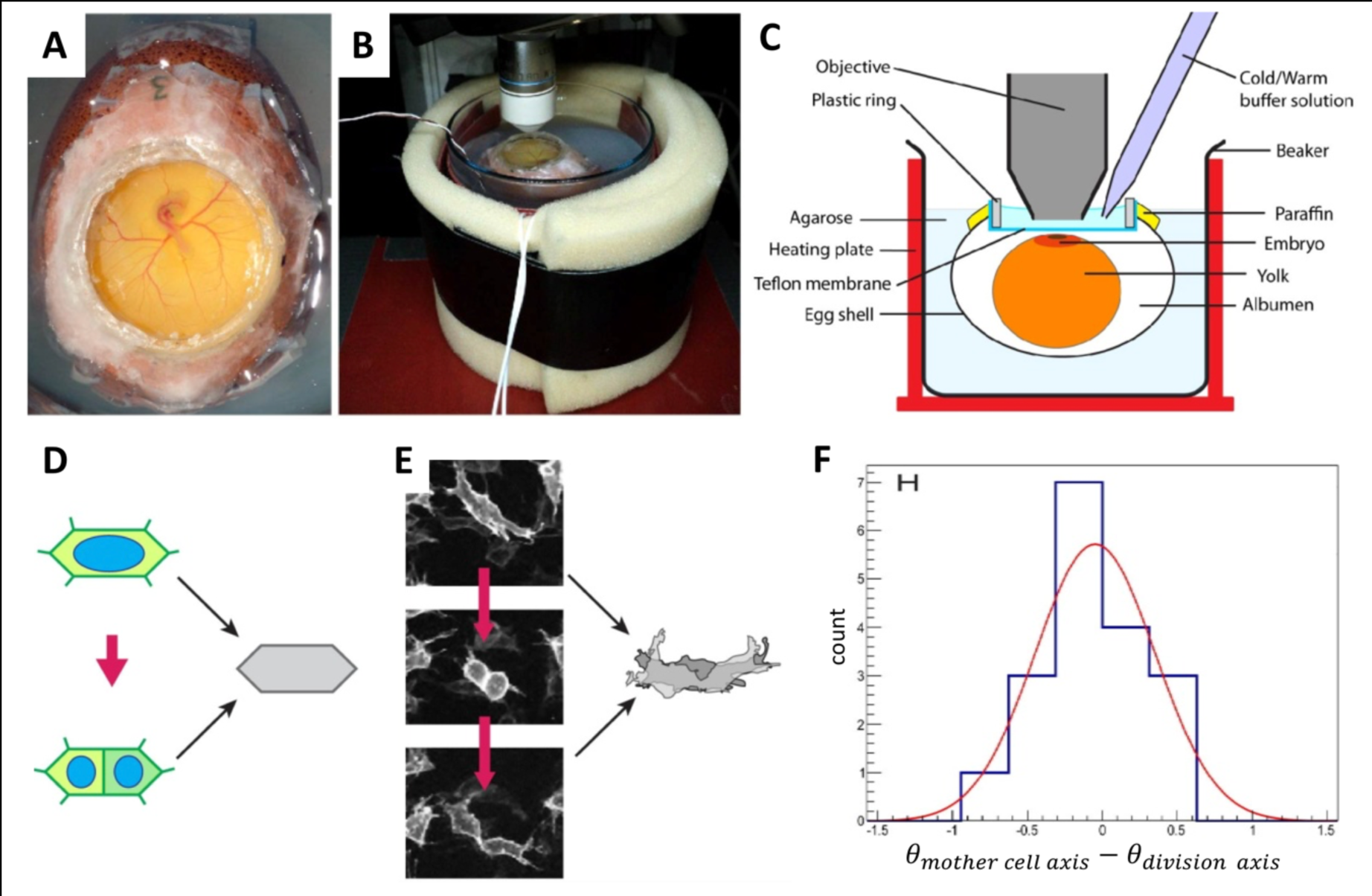
*In ovo* imaging of mesenchymal cell behaviour. (A-C) *In ovo* imaging technique. (A) Windowed chicken egg resealed with a ring covered by a Teflon membrane and embedded in agarose. (B) Beaker containing the egg, wrapped with a heating tape and insulation foam, and placed on a microscope stage adapted to increase clearance under the objective. (C) Applying cold buffer solution on the window stops the heart beat for imaging, while applying warm buffer solution maintains incubator-like conditions for normal development. (D) Hertwig’s rule: cell long axis determines division plane (E) Time-lapse images of cell division in the limb bud. As per Hertwig’s rule, mother cell shape predicts the daughter cells’ separation angle and orientation and matches the combined shape of the daughter cells (overlaid grey silhouettes). (F) Histogram of angle deviations between a mother cell’s longitudinal axis and its division axis (n=18 cells).

Throughout the entire duration of the time-lapse imaging, we could observe multiple cell divisions, a sign that our method did not significantly perturb normal embryonic development. Patterns of tissue development were normal, though somewhat slower than in an embryo maintained in an incubator. This was expected since chick embryos exposed to short periods of low temperature survive and hatch, but develop more slowly than those maintained at the optimal temperature (Suarez, Wilson et al. 1996). Embryos returned to the incubator after imaging continued to develop normally, providing the confidence that our technique gives access to dynamic images of healthy, normally growing tissue. The current study employed 18 time-lapse sequences covering up to 12 hours of development and these were analysed to characterize cell division, migration and convergence-extension movements.

### Cell Division Orientations Do Not Contribute to Limb Extension

In many tissues, cell-division orientation aligns with the direction of tissue elongation and is believed to promote anisotropic expansion (Baena-Lopez, Baonza et al. 2005, Tawk, Araya et al. 2007). We have previously shown that cell divisions in early limb buds are oriented toward the ectoderm, and thus in much of the bud divisions are almost perpendicular to the PD axis of limb bud elongation (Boehm, Westerberg et al. 2010). So, why does the limb bud mesenchyme not expand in the direction of cell division orientation?

We used our *in ovo* time-lapse imaging to monitor limb cells during their life cycle in detail. The size and shape of each mother cell is usually very similar to the combined territories of the daughter cells after separation (Figure 2D,E). Moreover, mesenchymal cells display a clear long-axis, and we found the relative positions of the two daughter cells after division to be highly correlated with the previous orientation of the mother cell (Figure 2F). Thus, mesenchymal cells appear to obey a version of Hertwig’s rule – the cell division orientation is largely determined by the long-axis of the mother cell (Figure 2D).

Cell-division orientation obeying Hertwig’s rule does not in itself suggest anisotropic growth (Gillies and Cabernard 2011). In the limb bud, because daughter cells and the mother cell occupy a similar region of space, cell divisions do not create anisotropic stresses on the tissue and DV-oriented cell divisions need not lead to a DV expansion of tissue. However, subsequent cell growth could lead the daughter cells to each replicate their mother cell’s shape and orientation. In the absence of cell reorganization, such growth could theoretically contribute to expansion along the DV axis. These observations therefore suggest that coordinated cell rearrangements may counteract DV expansion.

### Limb Cells Have Complex Shapes with Filopodia Protrusions Biased Toward the Ectoderm

Conflicting evidence and conclusions have been reported regarding the importance of active cell migration in limb elongation. Li & Muneoka reported that DiI labelled cells could migrate towards an implanted bead soaked in FGF (Li and Muneoka 1999). Since FGF is normally secreted from the distal-most ectoderm (the AER), this suggested that limb bud elongation might occur due to distal-ward cell migration (Schaller, Li et al. 2001). Others have also found signs of active cell migration in chick and mouse limb bud (Gros, Hu et al. 2010, Wyngaarden, Vogeli et al. 2010). However, cell orientation are perpendicular to the PD axis in large regions of the limb bud (dorsal and ventral regions) (Boehm, Westerberg et al. 2010, Gros, Hu et al. 2010).

To resolve these conflicting observations and statements we made use of the membrane-targeted GFP labelling to characterise the cell shapes in greater detail, including the fine projecting filopodia. We expect migrating cells to display a higher protrusive activity towards their direction of movement. Our results show that while most cells have a discernible longer primary axis, their shapes range from mono-, to bipolar, to complex (Figure 3A). The cells have an array of filopodial protrusions, which are up to 3 cell diameters long. Often, a structure similar to a small lamellipodium can be seen at the base of multiple filopodia. When observed over time, the filopodia are very dynamic, with an average of 0.46 filopodia being extended or retracted per cell, per hour (372 filopodia analysed from 8 limb buds) (Figure 3B, Table S2).

**Figure 3.**
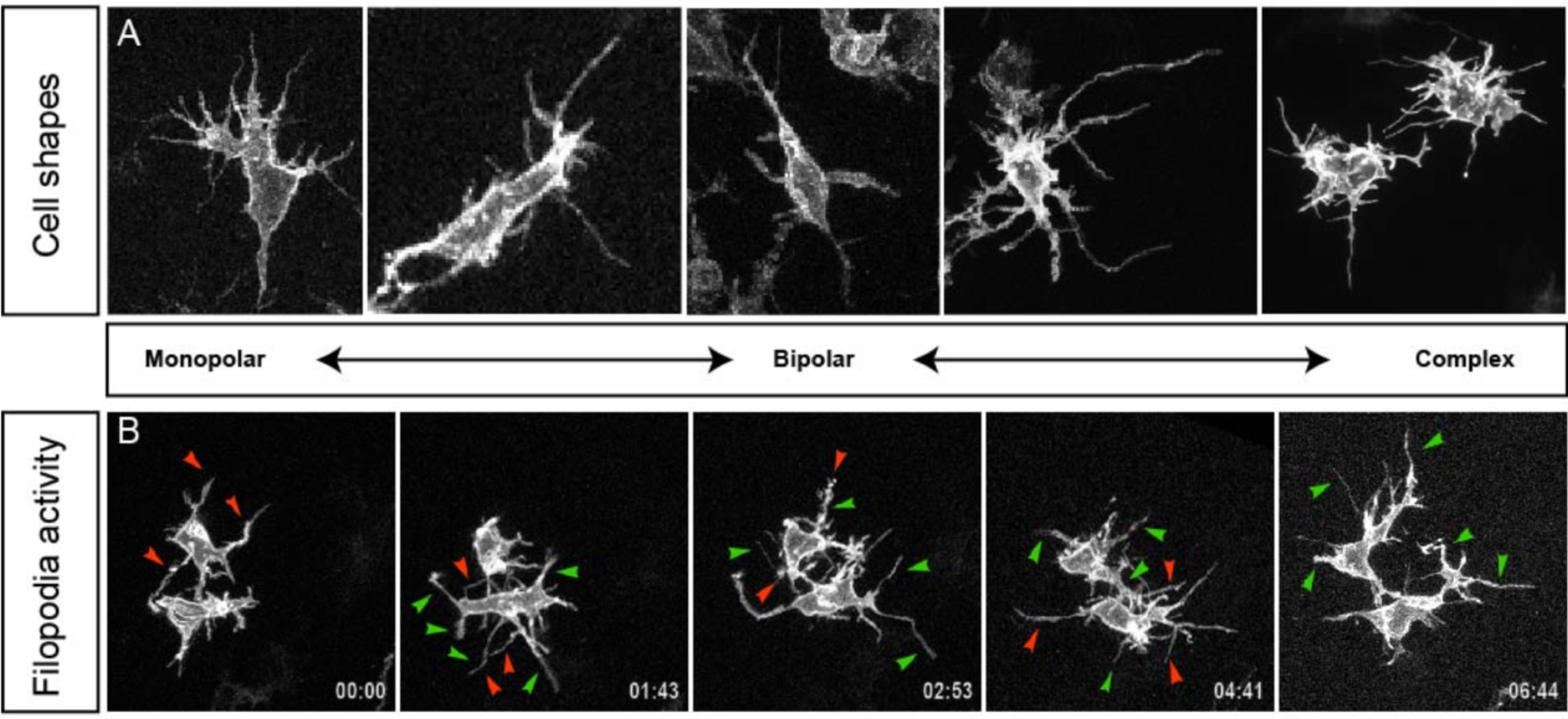
Cell shapes and filopodial activity. (A) Typical cell shapes in limb mesenchyme visualized with electroporated gpiEGFP. Cell shapes include monopolar, bipolar, and spider-like multipolar. (B) Time-lapse sequences showing filopodial activity, highlighting positions where filopodia will form (green arrow-heads) and filopodia that will retract (red arrow-heads).

### Active Cell Migration is Rare

The observed complexity of cell shapes emphasized the need for dynamic time-lapse analysis of individual cell displacement over time. An important question was whether limb bud elongation is driven primarily by active cell migration in the distal direction, as previously suggested (Li and Muneoka 1999). Our *in ovo* time-lapse data revealed that a few mesenchymal cells exhibit very clear migratory behaviour (Figure 4A,B) – actively crawling past their neighbours – but that they are a very small sub-set of mesenchymal cells, representing only 6% of cells at any instant (n=200) (Table S1). These cells are usually monopolar, with an active lamellipodium pointing in the direction of migration and a tapering trailing edge on the opposite side (Figure 4A,B). However, overall these migrating cells shows no clear bias in any particular direction. The only consistent observation is that after cell division daughter cells commonly migrate in opposite directions away from each other (Figure 4C,D).

**Figure 4.**
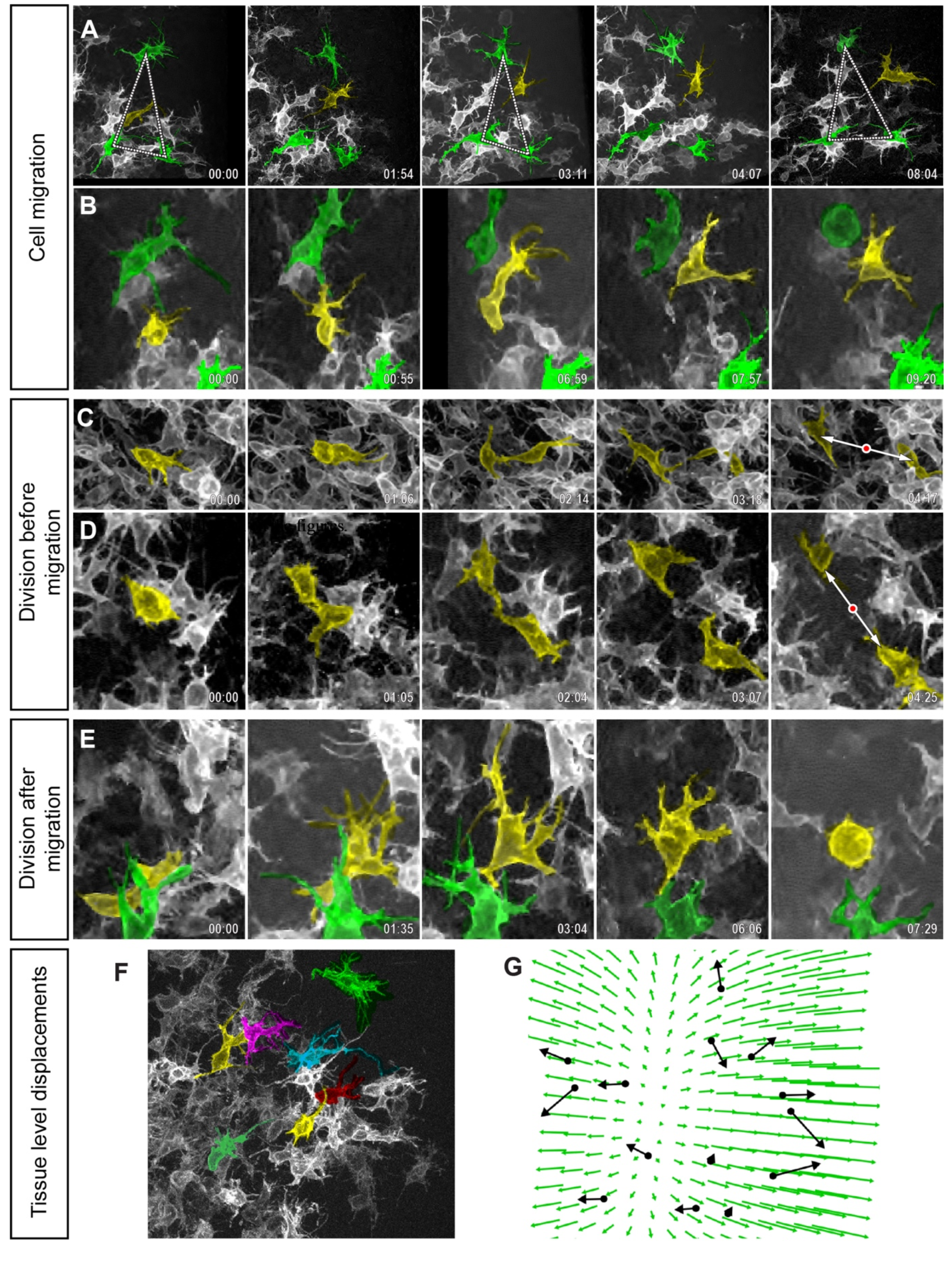
Analysis of relative cell movements. (A, B) Time-lapse sequences showing cell migration. A random sub-set of mesenchymal cells have been fluorescently labelled with gpiEGFP. A few labelled cells have been manually colour-coded to highlight their movements, The yellow cell is migrating; green cells and the triangle serve as reference points to highlight the relative displacement of the yellow cell. Apart from the yellow-labelled cell, no cells show significant movement relative to each other during the 8 hours of the movie. (C, D) After cell division, daughter cells often migrate in opposite directions (double-arrow). (E) In rare cases, cell division occurs after a cell has migrated. (A-E) In time-lapse series, distal is to the right and proximal is to the left. (F) Tracking a larger number of cells to deduce global tissue movements. (G) A bias can be seen in the cells moving apart from each (black arrows) other along the PD axis, which is described more generally by fitting a vector field (green arrows).

The lack of coordinated active migration argues against cell migration being important for limb bud elongation,but raises the question whether the migrating cells represent a distinct sub-population within the mesenchyme. To address this question, we assessed the persistence of migratory behaviour. In all cases observed, a migrating cell did not maintain this activity during the entire time-lapse sequence (which were typically 8-12 hours). Instead, during the course of the time-lapse relatively stationary cells occasionally start to migrate, or the reverse – migratory cells usually become stationary after a few hours. Before or after a migratory period, the cell was indistinguishable from its neighbours. We also assessed whether cells have a greater tendency to migrate at a certain point in the cell cycle, and found that migration events just after cell division were 10 times more common than just before cell division (48% of total migration events, versus 5%, for n=44 migrating cells, Figure 4C,D versus E). Taken together these data suggest that the mesenchymal tissue is not composed of two distinct populations (a migratory one, and a stationary one). Instead, the tissue appears as a relatively homogeneous population in which individual cells have the intrinsic ability to switch between two behaviours. Indeed, this switch has been observed previously – when plated *in vitro*, all mesenchymal cells have the ability to switch to a migratory state (Fisher and Solursh, 1979). By contrast, in their natural, *in vivo* context within the limb, the cells spend most of their time in a relatively stationary state, with active extension and retraction of filopodia, and only occasionally converting to a migratory behaviour, particularly after cell division.

### Cells separate apart along the PD axis

Since active cell migration is such a rare event, and therefore cannot be driving the bulk elongation of the limb bud, we sought evidence for an alternative mechanism. If the tissue were undergoing a form of convergent extension, then we should be able to observe mesenchymal cells significantly moving away from each other along the PD axis, compared to the other directions. In a classical convergent extension, with no tissue proliferation, cells would move towards each other along the converging axes (perpendicular to the PD axis). However, if the cells were rapidly proliferating then this overall tissue expansion could potentially compensate the convergence so that no strong movements would be seen along these axes – only extension along the PD axis would be significant.

To address this question, we took advantage of the salt-and-pepper labelling of mesenchymal cells follow electroporation, which allowed us to track distinct cells with confidence. From each time-lapse movie we could track up to 15 cells, and indeed a movement of cells away from each was most observable along the PD axis (black arrows in Fig.4X). To extract a more general measurement of the relative movements we calculated the vector field which best fits the individual cell movements (Fig.4X). This helps to visualise the strong divergence along the PD axis compared to the other axes, and also to estimate the growth rate, which was observed as 21% increase in area over 10 hours, which is equivalent to a 3D volumetric growth of 33%.

In principle, compression along the DV and AP axes could be extrinsic to the mesenchymal cells (for example, a squeezing force applied by the ectoderm), or intrinsic to the mesenchyme (if the cells actively intercalate between each other along the DV/AP directions). All available evidence supports the second of these two scenarios. If the ectoderm was mechanically important for squeezing the mesenchyme, then removing regions of it would have a rapid dramatic impact on elongation. However, the opposite result has been reported (Saunders 1948, Martin and Lewis 1986, Boehm et al. 2010). Additionally, external compression forces would predict that cells become flattened in an orientation parallel to the ectoderm, as is indeed seen 3 days later by chondrocytes which are oriented perpendicularly to the compressive forces of elongating long bones (Ahrens, Li et al. 2009). By contrast, mesenchymal cells in the elongating limb bud show no such arrangement and are consistently oriented in the opposite direction (see the last section of this study). The highly dynamic extension and retraction of filopodia suggests that cells could indeed pull on each other, such that the compressive forces would be generated by the mesenchymal tissue itself.

### The Global Arrangement of Cellular Orientations

In the case of a convergent extension mechanism, cellular orientations need to be mostly perpendicular to the axis of limb elongation (Figure 1D), so we sought to map the arrangement of cellular orientations within the limb bud. We were able to analyse three types of orientation marker: The orientation of filopodia is a very direct measurement, and mechanistically relevant for cellular intercalation, but due to the fluorescent labelling is rather sparse. To create a more complete mapping we therefore chose to supplement the filopodial data with two other measurements of cellular orientation: the position of the Golgi body relative to the nucleus, and the orientation of cell division.

In most regions of the limb bud, a cellular orientation towards (or away from) the nearest ectoderm will be perpendicular to the PD axis. This is true everywhere except the distal mesenchyme where the nearest ectoderm will instead be parallel to the PD axis. We therefore assessed the relationship between filopodia orientation and the nearest ectoderm for a collection of well-labelled mesenchymal cells (Fig.5). The results showed that filopodia are indeed biased towards the nearest ectoderm (Fig.5B-F). Examining Golgi orientation found a similar result (Fig.5G,H). Since neither of these biases were 100%, we asked whether they were independent of each other. Fig.5I shows that the Golgi and filopodial biases correlate with each other, and Fig.5J that this correlation even holds when both measurements are pointing away from the ectoderm. As a result, cells whose Golgi are facing away from the ectoderm but whose filopodia are extending towards the ectoderm (and *vice versa*) are very rare.

The strong correlation between Golgi position and filopodial orientation suggests that they are reliable indicators of general cell orientation, so we mapped a much larger number of Golgi orientations across different regions of 3 transverse sections of stage HH23 buds (n=1221 cells). The maps confirm that the ectodermal bias holds true for dorsal, ventral and distal regions of the limb bud (Fig.5K). We also analysed these sections for cell division orientation and found a consistent bias towards the ectoderm in the dorsal and ventral regions (n=445 cells) (Fig.5L). By contrast, the distal region showed a much more random orientation of cell divisions. Together, these results confirm that the gross orientations of cell shapes and activities is perpendicular to the PD axis, except in the very distal regions.

### A Convergence-Extension Model for Limb Bud Elongation

Together these results point towards a model in which the limb bud mesenchyme is undergoing a convergent-extension processes, with cells intercalating along the AP-DV plane and extending the limb along the PD axis. But can such 3D arrangement of intercalations events lead to extension of the limb? Convergent-extension has often been modelled in 2D contexts (Weliky, Minsuk et al. 1991, Zajac, Jones et al. 2000, Zajac, Jones et al. 2003, Brodland 2006, Rauzi, Verant et al. 2008, Backes, Latterman et al. 2009, Vroomans, Hogeweg et al. 2015), but little investigation has been devoted to 3D geometries (Honda, Nagai et al. 2008, Belmonte, Swat et al. 2016). A recent 3D convergent-extension model demonstrated that when cells are bi-polarized and intercalate along a main axis, tissues converge along the axis of intercalation while expanding in the orthogonal plane (Figure 6A,B) (Belmonte, Swat et al. 2016). However, the limb requires a mechanism to convergence along two axes (DV and Anterior-Posterior, AP) while extending along the PD axis (Figures 1D, 6D). Here we propose that if locally all cells intercalate bi-directionally towards the nearest ectoderm, the net movement of the tissue will be along the PD axis (Figure 6C,D). The observed uniform proliferation of the limb mesenchyme observed by Boehm *et al*. (Boehm, Westerberg et al. 2010) might then counter-act the tissue convergence along the DV and AP axis while leaving the PD expansion unaffected (Figure 8).

**Figure 5.**
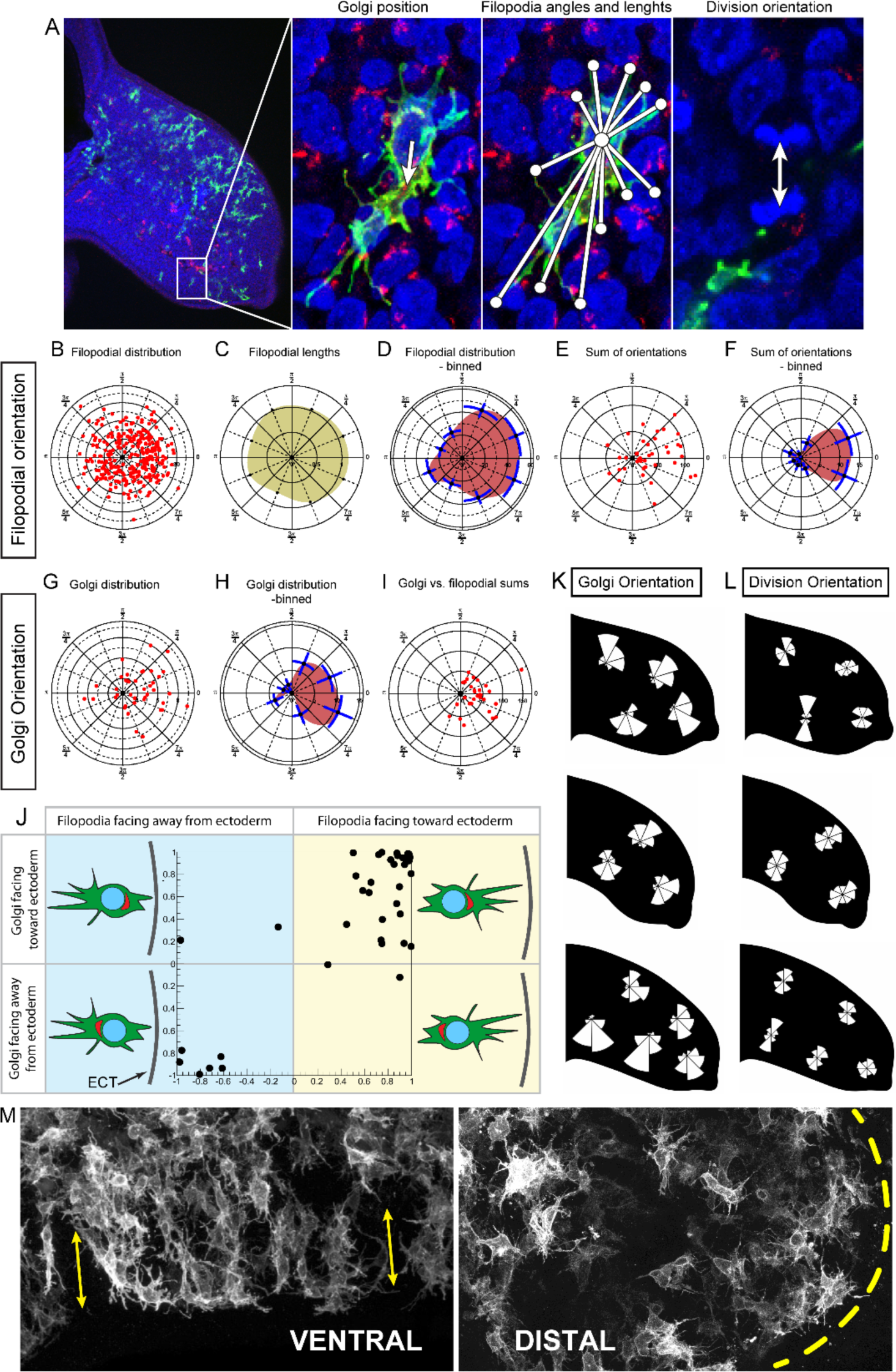
Global orientation of mesenchymal cells in the limb bud. (A) Markers of cellular orientation in HH21 stage limb buds. gpiEGFP labels the membrane of cells (green), thus revealing the shape and extent of filopodia, while the anti-GM130 antibody immunolabels the Golgi (red). Nuclei stained with DAPI (blue). Angles between nucleus and Golgi, nucleus and each filopodium tip, and division orientation were measured as shown, and summarised in the subsequent polar plots. (B) Polar dot-plot in which each red dot represents filopodium’s angle and length, with angle 0 (to the right) representing the direction to the closest ectoderm (n=40 cells). (C) Data as in (B), averaged over angular bins, reveals that the lengths of filopodia does not vary with angle. However, when binning the frequency of filopodia in different directions (D) a clear bias is revealed towards the nearest ectoderm. (E, F) If we analyse the data per cell (rather than per filopodium) then an even stronger bias is observed towards the ectoderm. This is seen either as the vectorial sums of filopodia for each cell (E), or the angular frequency distribution (F). (G) Polar dot-plot of the Golgi-to-nucleus distance, and an angular histogram of these data (H) show that most Golgis are oriented toward the ectoderm. (I) Since both filopodia (panels B-F) and Golgi (panels G,H) are oriented towards the ectoderm, we also plotted Golgi directly against filopodial orientation to confirm their direct correlation. (J) Plotting filopodial orientation against Golgi orientation shows that the majority of cells have both their filopodia and Golgi facing the ectoderm, a minority have both their filopodia and Golgi oriented away from the ectoderm, and almost no cells show a non-correlated arragment. (K, L) Spatial mapping of cell division and Golgi orientation (n=1221 for Golgi mapping, and n=445 for division orientation). Proximal tissue shows stronger orientation toward the nearest ectoderm for both measurements, while distal mesenchyme is organised for Golgi, but not for cell division orientation, which can be further seen in (M).

**Figure 6.**
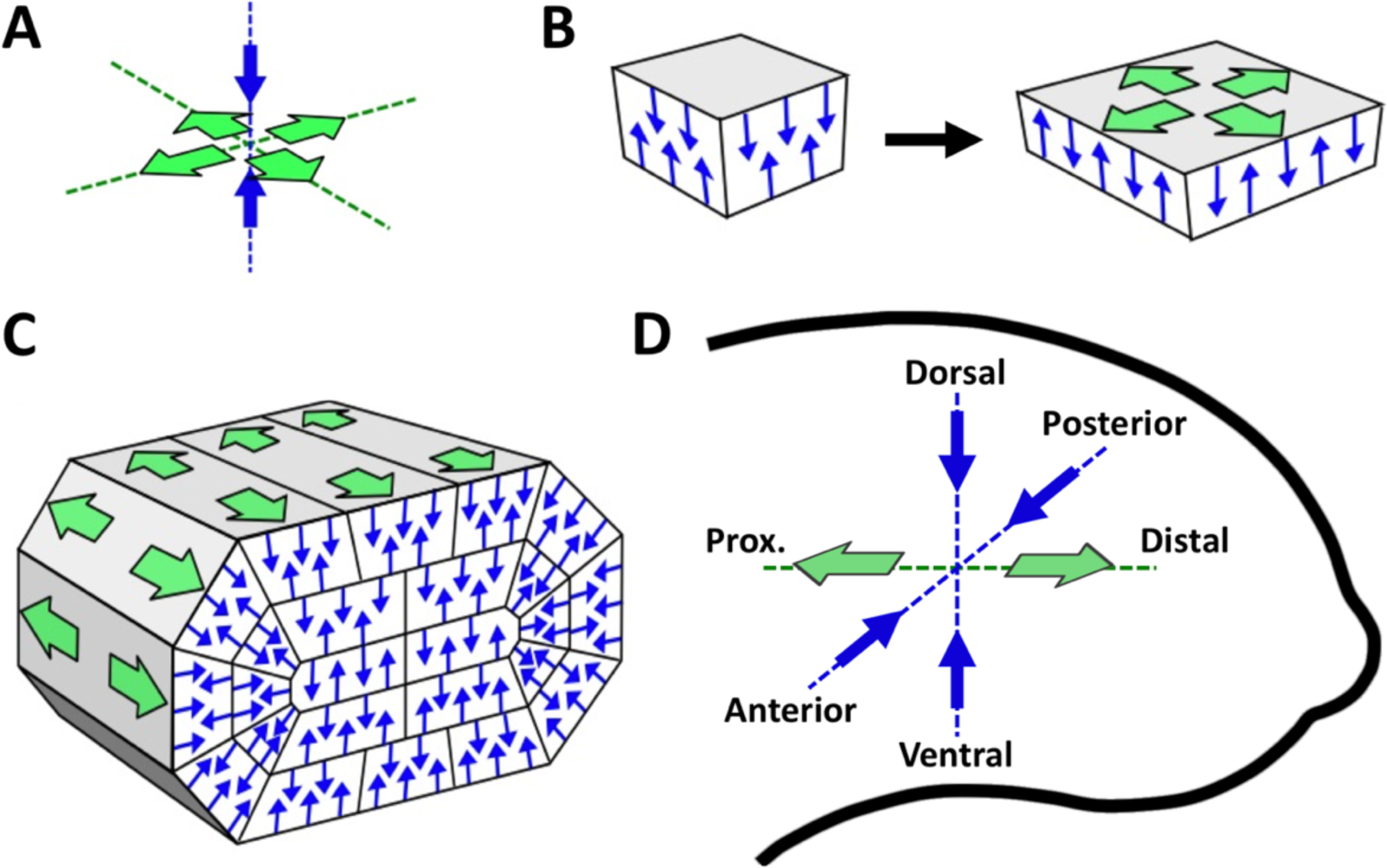
Convergent-extension model in the 3D limb bud. (A) Cell intercalation along a single-axis (blue arrows), creates an outward pressure in the orthogonal plane (green arrows). (B) When all cells within a tissue segment intercalate along the same axis, the tissue segment converges in the direction orthogonal to the axis and expands along the plane perpendicular to it. (C) Intercalation of cells towards/away the nearest ectoderm (blue lines) cancels extension along the directions tangential to the limb cross-section leaving only one axis of extension (green arrows). (D) Net convergence (blue arrows) and net extension (green arrows) movements in the developing limb bud leading to limb outgrowth.

**Figure 7.**
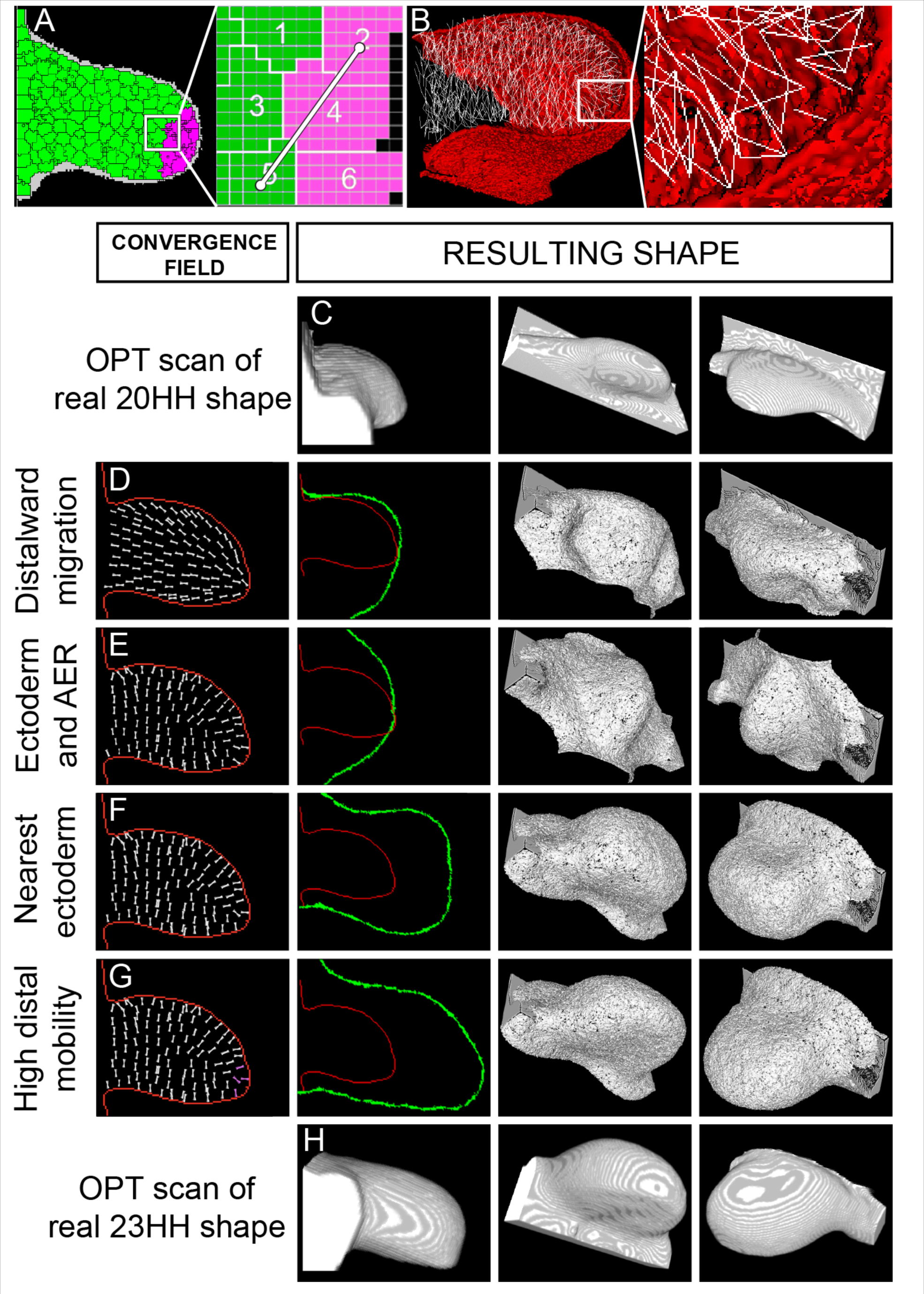
3D virtual-tissue simulations of the growing limb bud for mesenchymal cells with different contractile activities. (A) Left: DV x PD section of simulated limb (green – proximal mesenchyme; pink – distal mesenchyme; grey – ectoderm; black lines – cell boundaries). Right: To simulate contractile cell behaviour, pair of cells form transient contractile links (white bar) between their centers (white bullets). (B) 3D view of cellular links (white) from cells near the midplane in an AP x PD cutaway view of the simulated limb bud (Red – epithelium; transparent – mesenchyme). (C) Stage 20HH limb-bud acquired by OPT, used as initial state of the computer simulations. 3D views match those of the simulations in the same column. (D-G) Left column: midplane DV x PD cross-section of the initial limb bud shape in (C) (in red), and orientation of preferred intercalatory directions (white lines). Second column, same midplane cross-section of the initial (red) and final (green) simulated limb-bud shape. Rightmost columns show two perspective views of final (stage 23 HH) simulated limb-bud shapes in 3D. Row (D) Contractile links extend towards the nearest point of AER. (E) Orientation of the contractile links is influenced both by the AER and the nearest ectoderm. (F) Contractile links extend preferentially towards (and away from) the nearest point in the ectoderm. (G) Same as in (F), but filopodia in the distal region (pink) have shorter lifetimes than those in the proximal region. (H) Stage 23HH limb-bud acquired by OPT in views matching those of the simulations in the same column. Simulations begin from the shape of the stage 20HH limb bud in (C). In all simulations cells grow isotopically at a uniform rate and divide along random axes.

**Figure 8.**
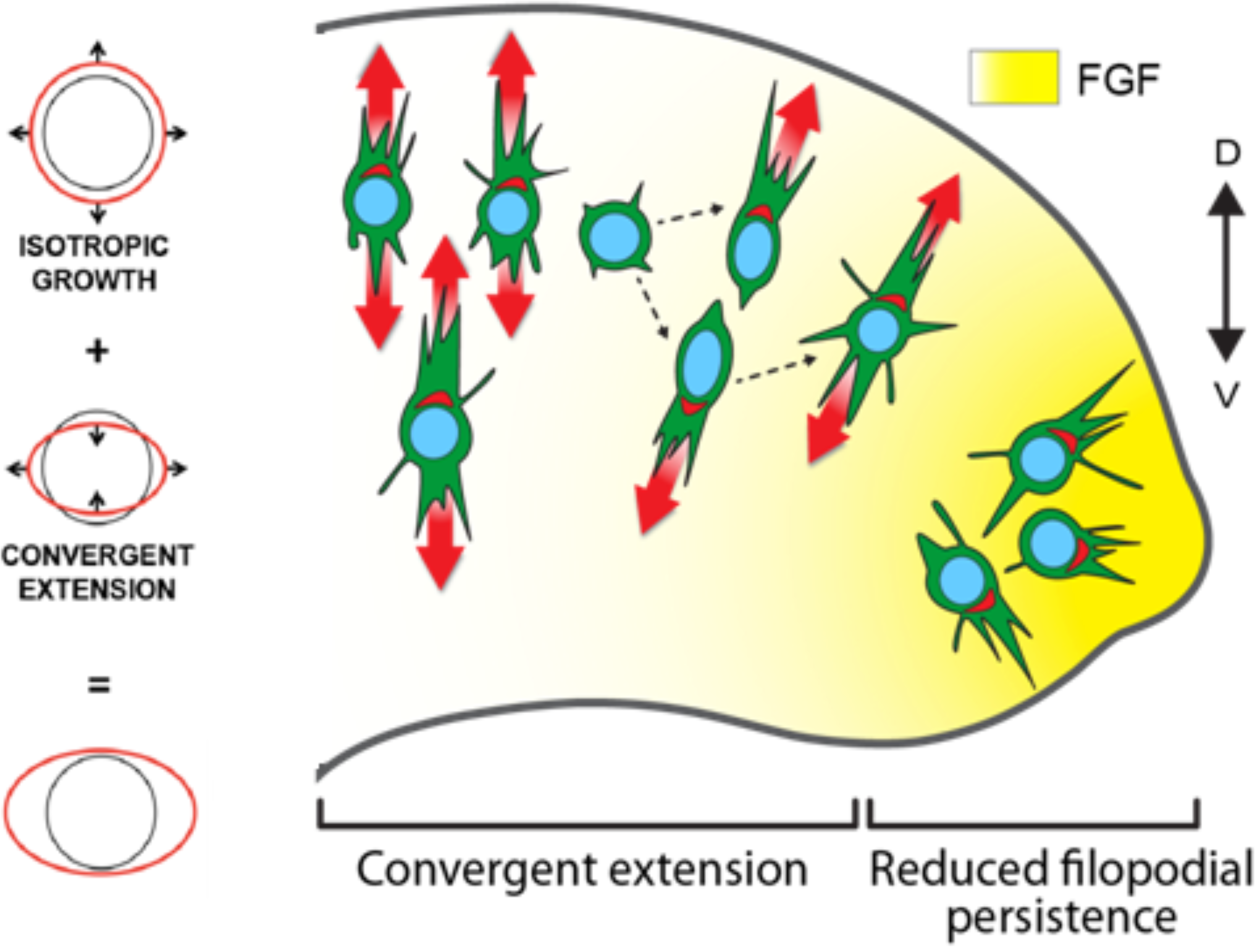
Proposed model of early limb bud morphogenesis. In most of the limb bud, cells are oriented perpendicular to the PD axis, and perform intercalatory movements (red arrows). Occasionally they divide, which involves briefly rounding-up into a spherical shape, then splitting into 2 daughter cells which initially take up a similar combined shape as the original mother cell, and then migrate off in opposite directions. The daughter cell which migrated away from the nearest ectoderm then appears to re-orient its Golgi, since the majority of cells analysed have their Golgi closest to the ectoderm (see Fig.5). Finally, cells in the distal tip show greater motility, and therefore more fluid tissue behaviour, possibly due to faster filopodial activity under the influence of FGF signalling. The correct morphogenesis is thus achieved by a combination of convergent-extension in most of the bud, and a softer tip region whose lack of tension permits the expansion.

### A 3D Computational Model of Limb Bud Elongation

Having defined the mesenchymal activities at the local level, we next wanted to explore theoretically whether they could be combined at an organ level to produce the correct 3D shape changes. Using the open-source CompuCell3D (*CC3D*) multi-scale simulation environment (Swat, Thomas et al. 2012), we developed a set of virtual-tissue simulations to test a series of hypotheses for the underlying cell behaviours responsible for limb-bud elongation, ranging from pure distally-oriented cell migration, as proposed by (Li and Muneoka 1999), to pure AP x DV intercalation as our experimental results seems to suggest. We used a novel 3D cell intercalation model implemented in CC3D which allows the modelled cells to form links (or virtual protrusions), which pull on other cells along a preferred direction (Belmonte, Swat et al. 2016). For simplicity, the modelled filopodial protrusions generate pulling forces between the centres of the interacting cells (Figure 7A). At this point, it is unclear if the pulling forces of the cells are generated predominantly by the filopodia, lamellipodia, by the rest of their membrane or all of them. To compromise, we set the lengths of the links to be between 1 and 2 cell diameters. The rate of link creation, persistence, and bias in orientation were controlled by the parameters of the model (Belmonte, Swat et al. 2016).

We considered two sources of orientation information – the ectoderm as a whole and the AER. The former is defined a thin layer of cells maintained around the mesenchymal cells (Figure 7B, red tissue), and the virtual AER was defined as a thin, distal strip of ectoderm with its AP extent controlled to reflect the pattern of FGF8 expression (see Supplemental Experimental Procedures for more details). Since this study focuses on the biomechanics of limb-bud elongation rather than on the molecular control of cell behaviours, we did not explicitly simulate the various chemical gradients hypothesized to provide orientation to the mesenchymal cells. Instead, each cell has 2 vectors measuring the directions to the nearest points in the ectoderm and the AER. All simulations were performed with a fixed uniform proliferation rate, which is a close approximation to the measured distribution of growth rates (Boehm, Westerberg et al. 2010). A 3D optical projection tomography (OPT) reconstruction of a stage 20HH (3 day) chick limb bud defined the initial virtual-tissue geometry (Figure 7C) (Sharpe, Ahlgren et al. 2002). We validated the simulations by comparing the virtual limb shapes to an OPT reconstruction of a stage 23HH (3.5-4 day) limb bud.

Our first test was pure distally-oriented filopodia (i.e. all oriented towards the closest part of the AER). This represents the hypothetical scenario in which all cells are migrating distally by extending filopodia towards their distal neighbours and then attempting to pull themselves forward (Figure 7D). In this model, the limb bud fails to extend because although each cell attempts to crawl distally it does so by pulling its neighbour backwards. The net result of these PD-oriented contraction forces tends to shrink the bud along the PD axis and extend it in the DV x AP plane. In conjunction with the underlying growth of the virtual tissue, the mechanism produces a round ball of tissue instead of an elongated paddle like shape.

Next we tested the involvement of cell intercalation (in other words the same contractile links as above, but not parallel to the PD axis). Based on the cellular measurements described previously (Fig.5), which suggest that cells are oriented towards the closet ectoderm, we allowed the virtual ectodermal gradient to also influence the orientation of the contractile links. Interestingly, when both virtual gradients influence the orientation of intercalation, a realistic elongation of the limb bud was still missing, even when the AER influence was weak compared to the ectodermal influence (Fig. 7E, numerical details in Supplementary Information). Only a model based on the ectodermal signal alone gave a realistic degree of convergent-extension (Fig.7F), as the contractile links were mostly perpendicular to the PD axis in this case. Nevertheless, the resulting shape of this simulation did not reflect normal limb bud morphogenesis – after an initial phase of elongation the distal end of the virtual bud began to bulge along the DV axis, and elongation was reduced (compare Fig.7F and H).

### A new morphogenetic role for FGF signalling in the distal limb bud

It appeared possible that the morphogenetic abnormality of the previous simulation (Fig.7F) derived from incorrect cell activities in the distal tip. In our initial simulations all mesenchymal cells were behaving the same, neglecting any differences in cell behaviour in proximal and distal mesenchyme. Orienting intercalatory activities along the cell-ectoderm axis produces an AP and DV oriented contractile force in the proximal limb bud that causes limb elongation, but it causes a PD oriented contractile force at the distal tip of the limb bud, which opposes the extension. This could explain why the simulated limb shapes had a flattened tip. In comparison to the real limb buds, the difference could be that the tip mesenchyme is under the influence of the FGF signalling, which was implicated in stimulating cell motility (Reiter and Solursh 1982, Li and Muneoka 1999, Gros, Hu et al. 2010). Interestingly, FGFR1 mutant mice also exhibit limb buds that are shorter in the PD direction and broader in DV and AP (Verheyden, Lewandoski et al. 2005), similar to the simulation result shown in Figure 7F. In other contexts FGFR1 signalling has been shown to inhibit E-cadherin expression, thereby allowing cells to become more motile (Ciruna and Rossant 2001).

To test the influence of higher cell motility on the limb shape we tested a final new model in which we defined a distal zone of mesenchyme, based on its distance from the AER, and reduced filopodial persistence in that region (pink arrows in Figure 7G). The higher turn-over of links is equivalent to faster, more dynamic cell activity. This change to the computer simulation made a significant difference to the shape – the limb bud extended further distally, avoiding the bulging shape of the previous model, and matched the 3D geometry of a HH23 limb bud remarkably well (Figure 7H).

The model thus predicts that correct morphogenesis is possible when the majority of the limb bud mesenchyme shows ectodermally-oriented cell intercalation, and when these coherent contraction forces are reduced in the distal tip. The tip cells in our simulations must have shorter link life-times than the rest of the tissue, which in real limbs, could correspond to higher cellular motility induced by FGF signalling and our own observations of differences in cell shapes between distal cells and the rest of limb mesenchyme (Figure 5M).

## Discussion

Combining a novel *in ovo* two-photon imaging, 3D tomographic scanning and a multi-cell virtual-tissue simulation, we propose a new view of how events at the cellular level determine limb bud morphogenesis at the organ level. The measured orientations of cell divisions are not compatible with an important role in regulating tissue shape. Similarly, active migration is seen in a small number of cells (6%) and is neither coordinated nor biased along the PD axis. Instead we propose that the major driving force for tissue elongation is cellular intercalation, which leads to a process of growth-compensated convergent-extension (Figure 8). Using a 3D cell-based computer model to explore hypothetical scenarios for the spatial distribution of orientations, we obtained a clear theoretical prediction that active cellular intercalation towards the closest ectoderm is a viable model for limb bud elongation. Our map of Golgi and filopodial orientations shown that mesenchymal cells are indeed oriented in such direction.

Our model also proposes a new morphogenetic role for FGF signalling in the distal limb bud, where FGF increases random cell motility and in such way lowers cells’ ability to intercalate. Although throughout most of the bud, cellular intercalation is towards the nearest ectoderm and therefore perpendicular to the PD axis, in the distal tip ectodermally-oriented convergence would act in the wrong direction (parallel to the PD axis). The computer model shows that if distal cells intercalate less, this allows the correct shape changes to proceed. Unlike the rest of the limb bud, we observed that distal mesenchyme is less-tightly organized (Fig.5M). Additionally, higher FGF signalling correlated with faster cellular dynamics agrees with previous evidence that FGF signalling acts to stimulate random cell motility (Gros, Hu et al. 2010) However, at the tissue level our hypothesis differs greatly from the previously suggested mechanism of morphogenesis. Both in the tail bud and the limb bud the higher distal motility has been proposed to drive elongation through a process of “mass action” of randomly moving cells (Benazeraf, Francois et al. 2010, Gros, Hu et al. 2010). Here, by contrast, we propose that the primary driving force is convergent-extension in the proximal regions of the limb bud. The higher motility of the distal cells in our model renders the tip more fluid, thus preventing convergent-extension in this region, which would otherwise be oriented in such a way as to inhibit limb bud outgrowth.

It may be useful to clarify the apparent discrepancy between our conclusions, and previous reports of active cell migration (Gros, Hu et al. 2010, Wyngaarden, Vogeli et al. 2010). The movement of any cell can be explained as a combination of extrinsic and intrinsic forces, or large-scale movement and active cell movement. The first category describes the fact that every cell is pushed by the general flow of the tissue. This movement will generally be coherent, as for example, two cells near to each other in the distal region of the limb bud will both tend to move away from the body at a similar speed. However, this coherence says little about the active movements made by the individual cells. For example, if fluorescent micro-beads were distributed throughout the tissue and tracked over time, their movements would also be highly correlated, despite being passive objects that exert no active forces in the tissue. To understand what cells are actively doing (and therefore how the large-scale tissue movements are driven) requires examining their local activities. By subtracting the tissue movements from the cell movements, we saw that local relative cell movements are not very coherent – indeed cells move away from each other along the PD axis (Figure 4G) – which implied that cell intercalation might be a more plausible mechanism of limb elongation than distal-ward cell migration. Additionally, previous studies showed that the PD elongation of the tissue does not depend on mechanical forces from the overlying ectoderm (Saunders 1948, Martin and Lewis 1986), thus it is much more likely that the general distalward flow of tissue is driven directly by this observed local intercalatory behaviour.

Although we propose a different role for FGF signalling on limb morphogenesis, our analysis is compatible with previous proposals about which are the most important molecular pathways involved in limb morphogenesis (FGFs, Wnts and PCP). As mentioned above, FGFs promotes higher cell motility and lower adhesiveness (Verheyden, Lewandoski et al. 2005, Gros, Hu et al. 2010), which helps to lower the intercalation at the tip. Directionality is thought to be provided by Wnt signalling (Heisenberg, Tada et al. 2000, Kilian, Mansukoski et al. 2003, Gong, Mo et al. 2004, Gros, Serralbo et al. 2009), and Wnt5a has been the more obvious candidate, due to its involvement with non-canonical signalling and the PCP system (He, Xiong et al. 2008, Wang, Sinha et al. 2011). Our results open new questions about how the PCP pathway could be involved. The Wnt5a gradient is along the PD axis with highest concentrations distally. Cell orientations in much of the bud are essentially DV, but in the distal region they are PD. For Wnt5a to specify these orientations would require opposite responses under the AER compared to under the dorsal and ventral ectoderm. We would suggest that the orientation may rather be achieved by a general ectodermal signal, such as Wnt3a (Parr, Shea et al. 1993, Barrow, Thomas et al. 2003), so that – well as under the AER.

Regarding generality of the model beyond limb development, it is important to highlight that despite superficial similarities with a recent proposal for primary axis elongation (Mongera et al. 2018), the two models are significantly different. The “fluid-to-solid transition” model of Mongera et al. also involves a distal (posterior) region of mesenchyme that is more fluid than the more proximal (anterior) tissue. However, in that model the force generation for physical elongation comes primarily from active tissue growth in the fluid region. The solid region acts as a rigid base imposing mechanical constraints on the growing region – in particular, preventing its expansion laterally and anteriorly, and thus mechanically guiding it to expand in the posterior direction. By contrast, in our model of limb morphogenesis, it is the growth and intercalatory activity of the more rigid tissue, which generates the active forces for limb elongation. Fluidity in the distal limb bud is only necessary to avoid “blocking” elongation. It is intriguing that the spatial distributions of mechanical properties may be similar, despite a fundamentally different causal mechanism for elongation.

## Experimental procedures

We developed an *in ovo* live two-photon imaging technique to analyse cell behaviour, and used the Glazier-Graner-Hogeweg (GGH) computational model with Compucell3D interface (Swat, Thomas et al. 2012) to mathematically test different proposed mechanisms behind limb morphogenesis. We also performed limb bud section immunostaining and statistically analysed mesenchymal cell orientations at different stages. Additional details on techniques and procedures are listed in the Supplemental Information.

## Acknowledgments

The authors would like to thank Drs. Miguel Torres, Rico Coen, Sevan Hopyan, Miquel Marin-Riera and Xavier Diego for critical reading of the manuscript. JMB is the recipient of the QuanTissue Short Visit Grant 4892. GL-P and JS were funded by the CRG and MINECO, Plan Nacional grant (BFU2010-16428).

