## Supplemental Information for "Cellular mechanisms of chick limb bud morphogenesis"

### Table of contents

**TABLE OF CONTENTS** 1

**SUPPLEMENTAL FIGURES** 2
FIGURE S1. MESENCHYMAL CELL ORIENTATION ANALYSIS, RELATED TO FIGURE 5 2
FIGURE S2. CELL INTERCALATION MODEL. 3
FIGURE S3. AER REPRESENTATION. 3
**SUPPLEMENTAL TABLES** 4
TABLE S1. MESENCHYMAL CELL MIGRATION, RELATED TO FIGURE 2 4
TABLE S2. COMPARISON OF FILOPODIAL ACTIVITY BETWEEN NORMAL AND FGF-INHIBITED LIMB BUDS, RELATED TO RESULTS: A NEW MORPHOGENETIC ROLE FOR FGF SIGNALING IN THE DISTAL LIMB BUD 4
TABLE S3: LIST OF PARAMETERS USED IN THE SIMULATION. 5
TABLE S4: LIST OF PARAMETERS USED FOR THE CELL-CELL CONTACT ENERGIES. 5
**SUPPLEMENTAL VIDEOS** 6
VIDEO S1. TIME-LAPSE VIDEO OF CHICK LIMB BUD MESENCHYME, RELATED TO FIGURE 2 6
VIDEO S2. COMPUTER SIMULATION OF CELL MIGRATION DURING LIMB MORPHOGENESIS, RELATED TO FIGURE 4D 6
VIDEO S3. COMPUTER SIMULATION OF CELLS PULLING IN THE DIRECTION BETWEEN THE NEAREST ECTODERM AND THE AER DURING LIMB MORPHOGENESIS, RELATED TO FIGURE 4E 6
VIDEO S4. COMPUTER SIMULATION OF CELL INTERCALATION DURING LIMB MORPHOGENESIS, RELATED TO FIGURE 4F 6
VIDEO S5. COMPUTER SIMULATION OF CELL INTERCALATION WITH THE DISTAL TIP SOFTENING DURING LIMB MORPHOGENESIS, RELATED TO FIGURE 4G 6

**SUPPLEMENTAL EXPERIMENTAL PROCEDURES** 7

IN OVO ELECTROPORATION 7
IN OVO IMAGING 7
REGISTRATION OF IN OVO IMAGES 7
CELL TRACKING AND GROWTH ANISOTROPY ANALYSIS 7
IMMUNO-STAINING 7
CELLULAR POTTS/GLAZIER-GRANER-HOGEWEG (*CP/GGH*) COMPUTATIONAL MODEL 8

**SUPPLEMENTAL REFERENCES** 11

#### Supplemental figures

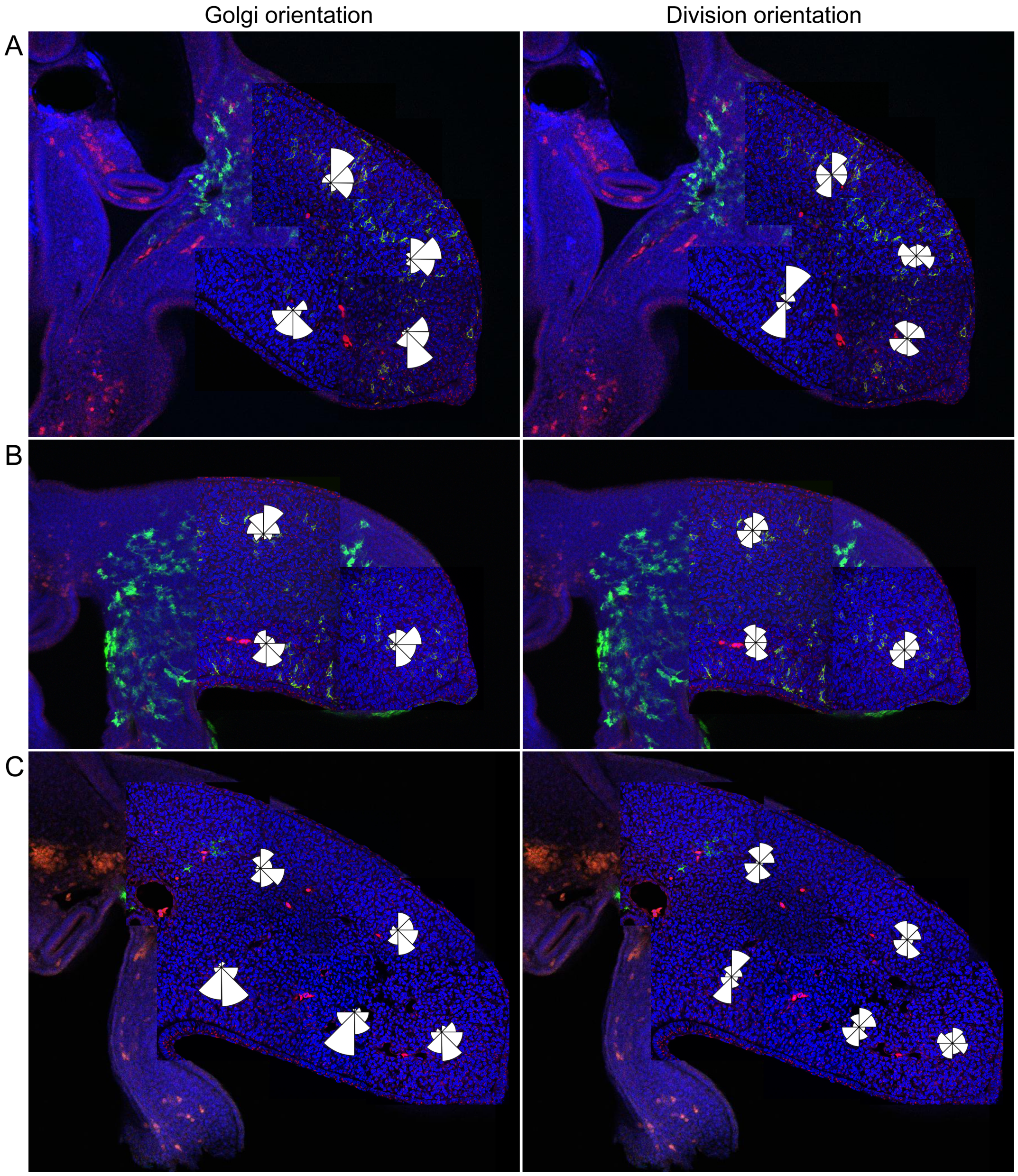


**Figure S1. Mesenchymal cell orientation analysis, Related to Figure 5**

(A-C) Confocal micrographs of 23HH stage limb buds used for Golgi orientation and cell division orientation assessment (see Figure 5J, K).

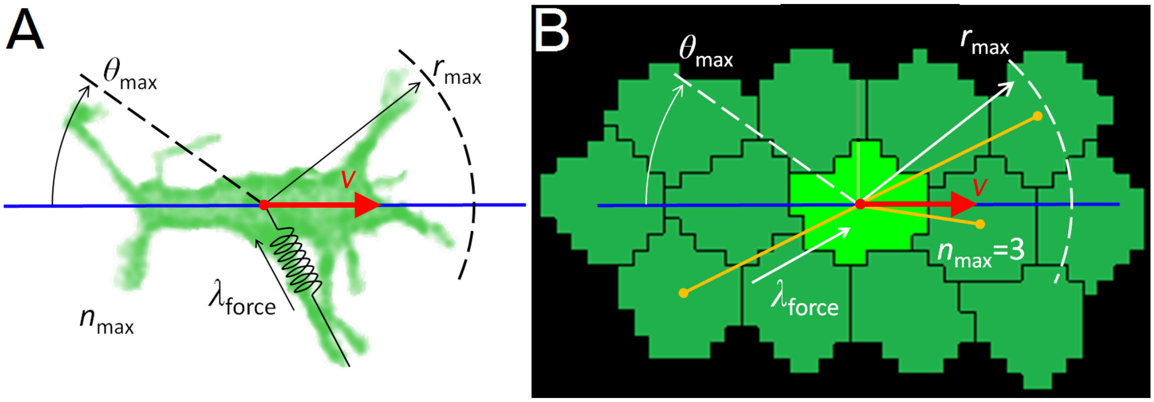


**Figure S2. Cell intercalation model.**

(**A**) Image of a bipolar cell in chicken limb-bud mesenchyme overlaid with model parameters. (**B**) Snapshot of a CP/GGH computer simulation of pulling force, overlaid with model parameters. Given an intercalation vector (in red), an intercalation axis is specified (x-direction) and the cell pulls a given set of neighboring cells ($n_{max}$) that lies inside a fixed interaction range ($r_{max}$) and within a critical angle ($\theta_{max}$). Yellow lines in (B) shows cell neighbor puller by the central cell. (Image adapted from Belmonte et al. 2016)

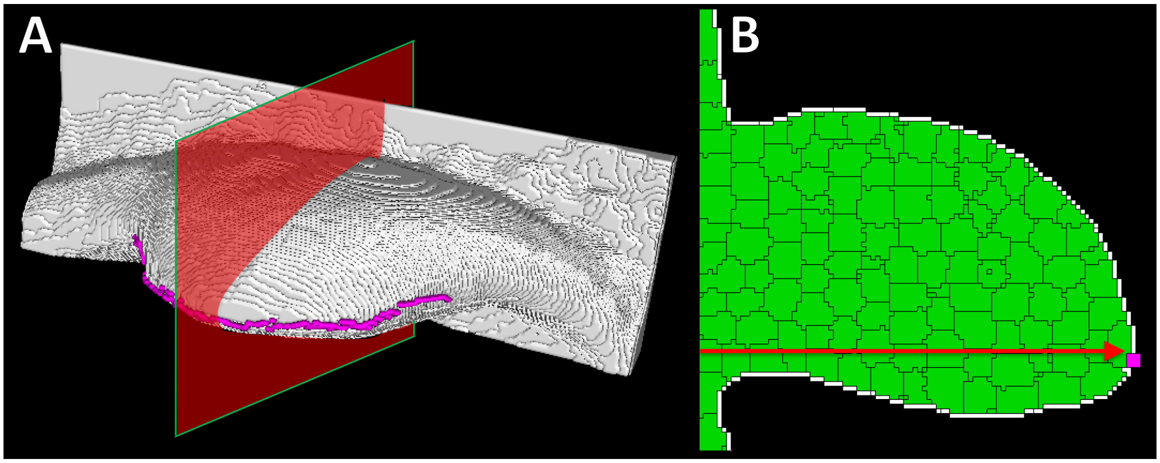


**Figure S3. AER representation.**

(**A**) 3D view of simulated limb bud (white) with AER location (magenta). Red panel show cross-section cut across the anterior-posterior axis. Dorsal side up. (**B**) 2D cross-section view of simulated limb-bud showing limb mesenchymal cells (green), ectoderm (white) and AER (magenta). AER at each plane is calculated as the most distal point of the ectoderm (red arrow).

#### Supplemental tables

| TL | Number of cells | Migrating cells | Non-migrating | Percentage of migrating | Time |
|---|---|---|---|---|---|
| **1** | 60 | 5 | 55 | 8.33% | 8h |
| **2** | 76 | 3 | 73 | 3.95% | 11h |
| **3** | 64 | 4 | 60 | 6.25% | 11h |
| **Sum** | 200 | 12 | 188 | 6% | |

**Table S1. Mesenchymal cell migration, Related to Figure 2**
Three time-lapse videos were analyzed for signs of cell migration. All together, 200 hundred cells were followed over a period of 8-10 hours and migrating cells were counted. We saw 12 cells to migrate for a short period of time, which corresponds to 6% of the whole cell population.

| **WT** | | |
|---|---|---|
| **TL** | **Extending** | **Retracting** |
| **1** | 6.79 | 7.21 |
| **2** | 4.85 | 6.08 |
| **3** | 4.67 | 5.35 |
| **4** | 4.00 | 3.50 |
| **Average** | 5.08 | 5.53 |
| **Average number of filopodia per cell** | 11.39 | |
| **Average of filopodial changes per hour** | **0.46** | |

| **SU5402** | | |
|---|---|---|
| **TL** | **Extending** | **Retracting** |
| **1** | 3.20 | 4.60 |
| **2** | 0.83 | 3.00 |
| **3** | 3.00 | 3.43 |
| **4** | 2.33 | 2.50 |
| **Average** | 2.34 | 3.38 |
| **Average number of filopodia per cell** | 8.91 | |
| **Average of filopodial changes per hour** | **0.32** | |

**Table S2. Comparison of filopodial activity between normal and Fgf-inhibited limb buds, Related to Results: A new morphogenetic role for FGF signaling in the distal limb bud**
Filopodial activity was compared between normal and SU5402-inhibited limb buds. We found that Fgf inhibition reduced the total number and the activity of filopodia. Normally, cells had an average of 11.39 filopodia per cell, with an average of 0.046 filopodial changes every hour. Fgf-inhibited mesenchymal cells had an average of 8.91 filopodia per cell, with only an average of 0.32 filopodial change every hour.

| Parameter | Symbol | Value (range) | Unit |
|---|---|---|---|
| Membrane fluctuation | $T_m$ | 100 | |
| Copy neighborhood order | $n_{copy}$ | 3 | |
| Cell target volume (M,DM) | $V_T$ | 216 - 432 | voxels |
| Volume constraint strength (M,DM) | $\lambda_V$ | 5 | |
| Growth rate (M,DM) | $\langle dV \rangle$ | 0.03 (0 - 0.06) | voxels/MCS |
| Cell target volume (E) | $V_{ET}$ | variable | voxels |
| Lambda Volume (E) | $\lambda_{EV}$ | 2 | |
| Division volume | $V_{max}$ | 432 | voxels |
| Target distance | $L_T$ | 3 | pixels |
| Pulling strength | $\lambda_P$ | 100 | |
| Maximum radius | $r_{max}$ | 2 | **cell** diameters |
| Maximum number of links | $n_{max}$ | 3 | |
| Maximum angle | $\vartheta_{max}$ | π/4 | radians |
| Interval between link formation (M) | $\tau$ | 50 | MCS |
| Interval between link formation (DM) | $\tau_{DM}$ | <u>15 - 32</u> | MCS |
| Contact neighborhood order | $n_{contact}$ | 5 | |

**Table S3: List of parameters used in the simulation.**

M = mesoderm; DM = distal mesoderm; E = ectoderm.

| **Cell types (σ)** | ***Medium*** | ***Mesoderm*** | ***Distal mesoderm*** | ***AER*** | ***Ectoderm*** |
|---|---|---|---|---|---|
| ***Medium*** | 0 | 30 | 30 | 10 | 8 |
| ***Mesoderm*** | | 10 | 10 | 10 | 12 |
| ***Distal mesoderm*** | | | 10 | 10 | 12 |
| ***AER*** | | | | 10 | 8 |
| ***Ectoderm*** | | | | | 6 |

**Table S4: List of parameters used for the cell-cell contact energies.**

Each value corresponds to the elements of the diagonally symmetrical contact energy matrix $J(\sigma_i, \sigma_j)$.

#### Supplemental videos

**Video S1. Time-lapse video of chick limb bud mesenchyme, Related to Figure 2**

Video of two-photon 40X magnification *in ovo* time-lapse imaging over a period of 11 hours. Cells were expressing membrane targeted gpi-EGFP to visualize the outline of the cells. Note the filopodial activity without major cell displacements.

**Video S2. Computer simulation of cell migration during limb morphogenesis, Related to Figure 4D**

Simulation of limb bud growth with cells pulling in proximo-distal direction as in case of distal-ward migration. See Figure 4D.

**Video S3. Computer simulation of cells pulling in the direction between the nearest ectoderm and the AER during limb morphogenesis, Related to Figure 4E**

Angle of cells was calculated to be between nearest ectoderm and the AER. See Figure 4D.

**Video S4. Computer simulation of cell intercalation during limb morphogenesis, Related to Figure 4F**

Simulation of cells pulling in the direction of the closest ectoderm.

**Video S5. Computer simulation of cell intercalation with the distal tip softening during limb morphogenesis, Related to Figure 4G**

Simulation of cells pulling in the direction of the closest ectoderm with cells in the distal region having decreased persistence of filopodia (in pink).

### Supplemental experimental procedures

**In ovo electroporation**

Fertilized eggs were purchased from a local farm and incubated on 38ºC for 65 h. In ovo electroporation was carried out as previously described (Suzuki and Ogura, 2008). Briefly, HH15 eggs were windowed, the vitelline membrane was removed, and 1 ml of 0.01% Fast green (F2752, Sigma Aldrich) in PBS was added on top of the embryo for better visualization. 1 µl of 1% Fast green was added to 5 µl of pCAGGS-gpiGFP vector (a kind gift from Fernando Giraldez Orgaz) at a concentration of 4 mg/ml. The DNA solution was injected into the embryonic cleome with microinjector (PicoPump PV820, WPI) and a sharp pulled glass pipette. A platinum anode was inserted beneath the embryonic endoderm and a platinum cathode was placed above the limb field. Three pulses of 4V, 60 ms pulse-on, 50 ms pulse-off were applied with CUY-21EDIT electroporator (Nepa Gene, Ichikawa, Japan) immediately after DNA injection. A small amount of 1% penicillin-streptomycin-amphotericin B (A5955, Sigma Aldrich) in PBS was added before the eggs were resealed and re-incubated for 24 h.

**In ovo imaging**

Time-lapse in ovo imaging technique was adapted from Kulesa and Fraser (Kulesa and Fraser, 2000). Embryos were first electroporated and after 24 hours of re-incubation (until they reached stage 21HH), the egg's window was covered with a ring with Teflon membrane attached to it (Fisher scientific, 13-298-83). The gap between the shell and the ring was sealed with paraffin and the egg was embedded in agarose in a beaker. The beaker was positioned under the microscope and wrapped in a heating plate to maintain the temperature at 38ºC. The developing limb bud was imaged with Leica SP5 upright two-photon confocal microscope with 40X/0.80 APO water immersion lens. 1µm z-stacks of the limb bud were taken every hour over periods of 8 to 11 hours. Typical depth was of about 90 µm. Before each time-point image was acquired, the heart was stopped by pouring ice cold PBS onto the egg. After the acquisition, the cold PBS was exchanged with warm PBS in order to raise the temperature back to 38ºC as quickly as possible. Between the acquisitions, PBS was removed to allow maximum oxygenation of the embryo through the Teflon membrane.

**Registration of in ovo images**

Due to the remaining heart-beat and shifting of the embryo during imaging, the obtained data needed post-processing in order to visualize the entire depth of the z-stack with maximum projection. Shifts between z sections were corrected by an in-house program for 2D post-registration in Matlab. Rotations between time points were manually corrected in Photoshop. Time-lapse videos were then analyzed in ImageJ.

**Cell tracking and growth anisotropy analysis**

Coordinates of cell positions at the initial and the last time point were measured in ImageJ and plotted as points on a graph. As a prediction of growth direction, a line was drawn from the center of the visualized mesenchyme to the nearest ectoderm. Marco

**Immuno-staining**

Embryos were dissected 24 or 48 hours after electroporation and fixed overnight in 4% paraformaldehyde in PBS at 4ºC, embedded in 5% low melting point agarose, and vibratome-sectioned at 200 µm. Sections were placed on Superfrost Plus glass slides and blocked overnight on room temperature (RT) in blocking solution: 10% heat-inactivated goat serum (G9023, Sigma Aldrich) in TBST (0.1% Tween 20 in 16TBS) with 0.01% Natrium Azide. Sections were incubated for 24 h on RT in blocking solution containing mouse anti-GM130 (1:250; 610822, BD Transduction Laboratories) and rabbit anti-GFP (1:500; 632460, Clontech). They were then rinsed in TBST with 0.01% Potassium azide overnight on RT and incubated in blocking solution with anti-mouse Alexa 568 (1:200; highly cross absorbed goat anti-mouse IgG antibodies, A11031, Invitrogen), anti-rabbit Alexa 488 (1:200; highly cross absorbed goat anti-rabbit IgG antibodies, A11034, Invitrogen), and DAPI (1:500; D8417, Sigma Aldrich) for 24 h on RT. Finally, slides were rinsed overnight on RT in TBST with 0.01% Potassium azide. Sections were scanned using Leica SP5 inverted confocal microscope with 40X/1.25 APO oil immersion and 63X/1.40–0.60 APO oil immersion objectives. Z-stacks with 1 µm depth were obtained.

Golgi, division and filopodia orientation was analyzed with ImageJ. For cell division analysis the angles between daughter chromatin centers during telophase or the angles perpendicular to the metaphase plate were measured.

**Cellular Potts/Glazier-Graner-Hogeweg (*CP/GGH*) computational model**

We implemented our 3D model of the chicken limb as a simulation using the Cellular Potts (CP), or Glazier-Graner-Hogeweg (GGH) model (Graner and Glazier, 1992) written using the open-source CompuCell3D software (Swat et al., 2012). The CP/GGH framework represents space as a regular lattice of sites, or voxels. Each simulated object (such as a cell, cell compartment, tissue or wall) is represented as a collection of voxels that forms a domain within this lattice. An effective energy ($H$) defines **cell**/**domain** properties such as size, mobility, adhesion preferences and distance constraints with other **cell**/**domains**. In our model **cells** have volumes, and interact via adhesion and dynamical cell-cell pulling forces, so that $H$ is given by three terms:

(Eq. 1) $$H = H_{Volume} + H_{Adhesion} + H_{Pulling},$$

The first term in the model's effective energy is a volume constraint to control the **cell**'s or **domain**'s size during the simulation:

(Eq. 2) $$H_{Volume} = \sum_{\sigma} \lambda_V (V(\sigma) - V_T(\sigma))^2,$$

where the sum is over all **cell**/**domains** ($\sigma$), $V(\sigma)$ is the current **cell**/**domain** volume, $V_T(\sigma)$ is the **cell**/**domain** target volume, and $\lambda_V$ is a lagrangian multiplier setting the strength of the constraint. **Cell** growth is modeled as a temporal evolution (increase) of the target volume parameter of the **cell** ($V_T(\sigma)$). Once the actual volume of a **cell** reaches a certain value ($V(\sigma) \geq V_{max}$), the **cell** is divided in half, at a random orientation.

Adhesion between **cells** is modeled with the standard Potts model internal energy term:

(Eq. 3) $$H_{Adhesion} = H_0 = \sum_{i,j} J(\sigma_{\boldsymbol{i}}, \sigma_{\boldsymbol{j}}),$$

where the sum is taken over all fourth-order neighboring pairs of grid coordinates $\boldsymbol{i}$ and $\boldsymbol{j}$; $\sigma_{\boldsymbol{i}}\ \sigma_{\boldsymbol{j}}$ are the **cell** domains at grid coordinates $\boldsymbol{i}$ and $\boldsymbol{j}$, respectively; and $J$ is the contact energy per unit area between those domains. Lattice sites belonging to the same generalized cell are assumed to have zero contact energy.

The last term in the effective energy is the pulling forces between cells. They are responsible for the **cell**-**cell** intercalation events that drive **tissue** extension, and were modelled after Belmonte et. al 2016. Each **cell** has an intercalation vector ($\boldsymbol{v}(\sigma)$, red arrow Fig. S2) that sets its axis of convergence (blue line in Fig. S2). Each intercalating **cell** selects all neighbors whose radial distance from it lies inside a given maximum range ($r_{max}$) and a given angle ($\vartheta_{max}$) with respect to the **cell**'s convergence axis (Fig. S1). From this pool the **cell** randomly selects all or a given number of neighbors to pull ($n_{max}$). The pulling is modeled with a spring force as shown below:

(Eq. 4) $$H_{Pulling} = \sum_{\sigma,\sigma'} \lambda_P (L(\sigma,\sigma') - L_T)^2,$$

where $L(\sigma,\sigma')$ is the actual distance between the two **cells**, $L_T$ is the target distance between them and $\lambda_I$is the strength of the interaction. After some relaxation time of $\tau$, all current links are discarded, and new ones are chosen based on the new neighborhood of each **cell**.

**Cell** dynamics in the GGH model provide a much simplified representation of cytoskeletally-driven cell motility using a stochastic modified Metropolis algorithm consisting of a series of index-copy attempts: the algorithm randomly selects a target site $\boldsymbol{i}$ and a neighboring source site $\boldsymbol{j}$; if different **cells** occupy those sites the algorithm sets ; $\sigma_{\boldsymbol{i}} = \sigma_{\boldsymbol{j}}$ with a probability given by the Boltzmann acceptance function:

(Eq. 5) $$P(\sigma_i \to \sigma_j) = \begin{cases} 1 & : \quad \Delta H \le 0 \\ e^{-\frac{\Delta H}{T_m}} & : \quad \Delta H > 0 \end{cases},$$

where $\Delta H$ is the change in the effective energy if the copy occurs, and $T_m$ is a global parameter controlling **cell** membrane fluctuations. A Monte Carlo Step (*MCS*) is defined as *N* index-copy attempts, where *N* is the number of sites in the cell lattice, and sets the natural unit of time in the computational model. The Metropolis algorithm evolves the cell-lattice configuration to simultaneously satisfy the constraints, to the extent to which they are compatible, with perfect damping (*i.e.*, average velocities are proportional to applied forces).

Our model of the limb has 3 **cell/domain** types, the mesoderm (M) which forms the bulk of the **limb** (green **cells** in Fig. 6A), the distal mesoderm (DM, on a subset of simulations) that lies close to the distal tip of the **limb** (magenta **cells** in Fig. 6A), and the ectoderm **cell** layer (E), that encloses the **limb** (white **cells** in Fig. 6A and red layer in Fig. 6B).

The geometry for the initial condition for the simulations was taken from OPT scans of 20 HH stage chicken embryos (Fig. 6C), where we segmented one of the fore limbs into a collections of lattice coordinates. We then subdivided the segmented limb into smaller units down to a 1:16 real cell to simulated **cell** ratio for the mesoderm. All lattice sites on the surface of the segmented limb were converted to **ectodermal cell** type and subdivided into smaller **cell** units.

Mesodermal **cells** of both types growth at an average rate of 0.03 pixels/MCS (at each MCS the actual increase in $V_T$ can be anywhere between 0 and 0.06 pixels/MCS), and divide at random division plane orientation once their volumes pass a certain value ($V_{max}$). **Mesodermal cells** close to the base of the simulated limb do not growth and have their shape frozen from the start of the simulation throughout. **Ectodermal cell** layer growth is dynamic and set as a function of its pressure ($\dot{V}_T = (V(\sigma) - V_T(\sigma))/2$), in order to just maintain the enclosure of the limb mesoderm.

We defined an AER region in the **ectoderm** by scanning the segmented limb along the anterior-posterior direction within the original width of the limb and finding the most distal **ectoderm** voxel at each plane (Fig S3). This is done periodically during the simulation to estimate the actual position of the AER. For the last model tested (the simulations in Fig. 6G), we define all mesodermal **cells** within 2 **cell** diameters from the AER as **distal mesodermal cells**. These cells have a reduced pulling persistence times compared to the mesodermal cells, with **cells** near the midline of the limb having the lowest persistence times.

For each **mesodermal cell** (M and DM) we calculate two vectors, the direction of the nearest ectodermal voxel ($\boldsymbol{v}_E$) and the direction of the nearest AER position ($\boldsymbol{v}_A$). We used these two vectors to calculate the **cell** intercalation vector $\boldsymbol{v}(\sigma)$, which will determine the individual axis of convergence of each **cell** within the simulated limb.

The cells for  the simulation shown in Fig. 6D have intercalation vectors pointed towards the AER ($\boldsymbol{v}(\sigma) = \boldsymbol{v}_A$); for the simulation shown in Fig. 6E cells have weighted intercalation vectors between the two directions ($\boldsymbol{v}(\sigma) = 0.7\boldsymbol{v}_E + 0.3\boldsymbol{v}_A$); and in Figs. 6F-G cells have intercalation vectors pointed towards the ectoderm ($\boldsymbol{v}(\sigma) = \boldsymbol{v}_E$). In the simulation shown in Fig. 6G cells all **mesoderm cells** from within 2 **cell** diameters from the AER are converted to **distal mesoderm**, and have a reduced pulling persistence time. All simulation parameters can be found in Tables S3 and S4.

Text for the main article

Experimental procedures

We developed an in ovo live 4D two-photon imaging technique to analyse cell behavior and used the Glazier-Graner-Hogeweg (GGH) computational model with Compucell3D interface to mathematically test different proposed mechanisms behind limb morphogenesis. We also performed limb bud section immunostaining and statistically analysed mesenchymal cell orientations at different stages. Additional details on techniques and procedures are listed in the Supplemental Experimental Procedures.
